# Alpha-mannosidase-2 modulates arbovirus infection in a pathogen- and *Wolbachia*-specific manner in *Aedes aegypti* mosquitoes

**DOI:** 10.1101/2022.03.18.484928

**Authors:** Nadya Urakova, Renuka E. Joseph, Allyn Huntsinger, Vanessa M. Macias, Matthew J. Jones, Leah T. Sigle, Ming Li, Omar S. Akbari, Zhiyong Xi, Konstantinos Lymperopoulos, Richard T Sayre, Elisabeth A. McGraw, Jason L. Rasgon

## Abstract

Multiple *Wolbachia* strains can block pathogen infection, replication, and/or transmission in *Aedes aegypti* mosquitoes under both laboratory and field conditions. However, *Wolbachia* effects on pathogens can be highly variable across systems and the factors governing this variability are not well understood. It is increasingly clear that the mosquito host is not a passive player in which *Wolbachia* governs pathogen transmission phenotypes; rather, the genetics of the host can significantly modulate *Wolbachia*-mediated pathogen blocking. Specifically, previous work linked variation in *Wolbachia* pathogen blocking to polymorphisms in the mosquito alpha-mannosidase 2 (αMan2) gene. Here we use CRISPR-Cas9 mutagenesis to functionally test this association. We developed αMan2 knockouts and examined effects on both *Wolbachia* and virus levels, using both dengue virus (DENV; *Flaviviridae*) and Mayaro virus (MAYV; *Togaviridae*). *Wolbachia* titers were significantly elevated in αMan2 knockout (KO) mosquitoes, but there were complex interactions with virus infection and replication. In *Wolbachia*-uninfected mosquitoes, the αMan2 KO mutation was associated with decreased DENV titers, but in a *Wolbachia*-infected background, the αMan2 KO mutation significantly modulated virus blocking. In contrast, the αMan2 KO mutation significantly increased MAYV replication in *Wolbachia*-uninfected mosquitoes and did not affect *Wolbachia*-mediated virus blocking. These results demonstrate that αMan2 modulates arbovirus infection in *Ae. aegypti* mosquitoes in a pathogen- and *Wolbachia*-specific manner, and that *Wolbachia*-mediated pathogen blocking is a complex phenotype dependent on the mosquito host genotype and the pathogen. These results have significant impact for the design and use of *Wolbachia*-based strategies to control vector-borne pathogens.

## Introduction

Dengue virus (DENV) (genus *Flavivirus*, family *Flaviviridae*) is an important human pathogen that is transmitted primarily by *Aedes aegypti* mosquitoes [1]. Mayaro virus (MAYV) (genus *Alphavirus*, family *Togaviridae*) is an emerging human pathogen that is transmitted mainly by *Haemagogus janthinomys* mosquitoes; however, *Ae. aegypti* mosquitoes are also competent vectors for this virus [2]. There are no approved vaccines or specific antivirals to prevent and manage disease outbreaks that are caused by either virus and thus novel strategies for disease control are needed to combat arbovirus infections. The use of the intracellular invertebrate-specific bacterium *Wolbachia* as a biological control agent against *Ae. aegypti* has emerged as an innovative vector control strategy to reduce arbovirus transmission. *Wolbachia* is useful because, when incorporated into *Ae. aegypti* mosquitoes, it suppresses vector populations via a reproductive manipulation called cytoplasmic incompatibility (CI) [3–5] and also prevents replication of viruses inside mosquitoes, a trait known as pathogen blocking (PB), thereby limiting subsequent virus transmission to humans [5].

*Wolbachia*-mediated pathogen blocking (PB) phenotypes in mosquitoes depend not just on the infecting *Wolbachia* strain, but also on many other factors including pathogen, infection type (natural vs. artificial), environmental conditions, and, importantly, host genetics [6–7]. Ford et al. found enough standing genetic variation in Australian *Ae. aegypti* to select for significant weakening of PB within a few generations of artificial selection, suggesting that the host genetic background can have a strong effect on PB [6]. Identified candidate mosquito host genes for this modulation were not the canonical suspects of mosquito innate immunity or detoxification; rather, they were primarily related to cell adhesion, Notch signaling, and cell cycle [6,7], highlighting our current lack of mechanistic understanding of the PB phenomenon.

Ford et al. identified single nucleotide polymorphisms in the non-coding region of the alpha-mannosidase 2 (αMan2) gene that were strongly associated with PB strength in *Wolbachia*-infected *Ae. aegypti* mosquitoes selected for high vs. low *Wolbachia*-mediated PB of DENV [6]. αMan2 is putatively involved in protein glycosylation [8], and thus could alter PB by modulating viral glycosylation. Protein glycosylation, the enzymatic attachment of oligosaccharide structures to the peptide backbone, is an important post-translational modification for both host cell and viral proteins [9–11]. In eukaryotic cells, glycosylation is responsible for many functions, including proper protein folding, trafficking, stability, receptor-ligand recognition, and cell adhesion [9]. Viruses do not have their own protein glycosylation machinery and employ host cellular enzymes for this purpose [10]. Glycosylation of viral proteins plays a crucial role in the lifecycles of many viruses, influencing virus infectivity, pathogenicity, and host immune responses [11,12]. Enzymes involved in protein glycosylation are important potential targets to control viral replication in eukaryotic cells [13,14]. However, how specific genes in these pathways affect arboviral replication in mosquito vectors is poorly understood.

In this study we used CRISPR-Cas9 gene editing to ablate the αMan2 gene in *Ae. aegypti* and examined effects of gene knock-out (KO) on mosquito vector competence for DENV and MAYV in both *Wolbachia*-infected and uninfected mosquitoes. Results demonstrated complicated interactions between gene KO, *Wolbachia* infection, and viral pathogen, highlighting the complex nature of *Wolbachia* PB phenotypes.

## Materials and methods

### Cells

African green monkey kidney (Vero, ATCC CCL-81) cells were obtained from ATCC (Manassas, VA, USA) and maintained in Dulbecco’s Modified Eagle Medium (DMEM) (Gibco/Thermo Fisher, Waltham, MA, USA) supplemented with 10% fetal bovine serum (FBS) (Gibco/Thermo Fisher), 100 ug/mL of streptomycin (Gibco/Thermo Fisher) and 100 units/mL of penicillin (Gibco/Thermo Fisher) at 37°C in 5% CO_2_. *Aedes albopictus* cells (C6/36) were obtained from Sigma-Aldrich, St. Louis, MO, USA, and maintained in RPMI 1640 medium (Gibco/Thermo Fisher) supplemented with 10% FBS (Gibco/Thermo Fisher), 100 ug/mL of streptomycin (Gibco/Thermo Fisher) and 100 units/mL of penicillin (Gibco/Thermo Fisher) at 28°C.

### Viruses

MAYV strain BEAN343102 (GenBank: KP842802.1) was obtained from BEI Resources, NIAID, NIH (Manassas, VA, USA). To produce MAYV stocks, virus was propagated on Vero cells for 24 hours and stored at - 80°C. DENV serotype 2 strain JAM 1409 [15] was propagated on C6/36 cells for 7 days as previously described [16]. MAYV stocks were initially quantified by plaque assay, while DENV stocks were initially quantified by qPCR. In mosquito infection experiments, viruses were quantified by focus-forming assay (FFAs; see below for specific methods).

### Antibodies

Mouse monoclonal anti-alphavirus antibodies (G77L) (#MA5-18173) were obtained from Thermo Fisher and used in FFAs to detect MAYV at a dilution of 1:40 and incubated at 4 °C overnight. Mouse monoclonal anti-flavivirus group antigen antibodies, clone D1-4G2-4-15 (produced *in vitro*) (NR-50327) were obtained from BEI Resources, NIAID, NIH (Manassas, VA, USA). These antibodies were used in FFAs for the detection of DENV antigens at a dilution of 1:500 and incubated at 4 °C overnight. Goat anti-mouse IgG (H+L) highly cross-adsorbed secondary antibodies, Alexa Fluor 488 (A-11029) were purchased from Invitrogen. Secondary antibodies were used in FFAs at a dilution of 1:1,000 and incubated at a room temperature for at least 3 hours or at 4 °C overnight.

### Plaque assay for the quantification of MAYV stocks

For quantification of MAYV viral stocks, Vero cells were seeded in 6-well plates at a density of 5×10^5^ cells/well. Ten-fold serial dilutions of virus stocks were prepared in PBS and 200 uL of these dilutions were used for infections. Cells were infected for 1 hour at 37°C, infectious media removed, and cells covered with 1 mL of complete DMEM medium containing 0.5% agarose. Three days post-infection, cells were fixed with 4% formaldehyde (Sigma-Aldrich) in phosphate-buffered saline (PBS) (Gibco/Thermo Fisher) for 25 min, agarose covers were removed, and cells were stained for 5 min using aqueous solution containing 1% crystal violet and 20% ethanol to visualize plaques.

### qPCR for the quantification of DENV stocks

Viral RNA was purified using Direct-zol RNA kit (Zymo Research) according to the manufacturer’s instructions and used as template in qPCRs. All primer sequences are in Table 1. qPCRs were set up using TaqMan™ Fast Virus 1-Step Master Mix (Thermo Fisher) and run on an ABI 7500 Fast Real-time PCR System (Applied Biosystems/Thermo Fisher). The thermocycling conditions were as follows: 50 °C for 5 min; 95 °C for 20 s; 35 cycles of 95 °C for 3 s; 60 °C for 30 s; 72 °C for 1 s; and 40 °C for 10 s. Product was detected by measuring the fluorescence signal from the FAM reporter. A standard reference curve of known quantities of a DENV-2 genomic fragment was used for absolute quantification by qPCR. The DENV-2 genomic fragment was inserted into a plasmid and transformed into *E. coli* as previously described [16]. The linearized and purified fragment was serially diluted ranging from 10^7^-10^2^ copies and were used to create a standard curve of DENV amplification. The standard curve was run in duplicate on each 96-well plate, and the limits of detection were set at 10^2^ copies.

**Table 1.**
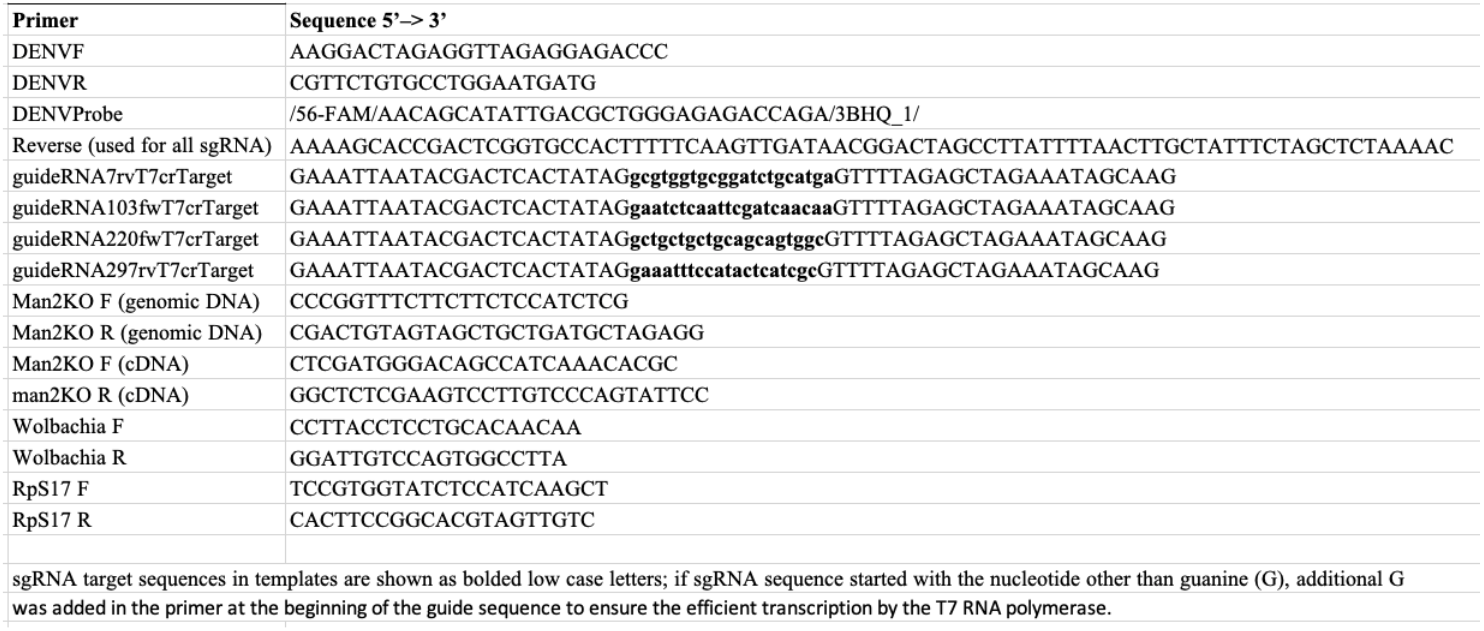
Primer and oligonucleotide sequences used in this study.

### Mosquito rearing

*Ae. aegypti* mosquitoes (Liverpool genetic background) expressing Cas9 protein in the germline (AAEL006511-Cas9; [17]) were provided by Dr. Omar Akbari, UC San Diego. *Ae. aegypti* mosquitoes stably infected with the wAlbB strain of *Wolbachia* were provided by Prof. Zhiyong Xi, Michigan State University. Mosquitoes were reared at the PSU Millennium Sciences Complex insectary under the following environmental conditions: 27±1°C, 12:12 hours light:dark diurnal cycle, 80% relative humidity. For reproduction, mosquitoes were maintained on expired anonymous human blood using a 37 °C water-jacketed membrane feeder. Larvae were fed on koi fish pellets (TetraPond). Adult mosquitoes were maintained on 10% sucrose solution.

### Preparation of single guide RNAs (sgRNAs)

The αMan2 Entrez Gene ID 5564678 gene sequence was used as a reference to design sgRNAs using CRISPOR [18]. sgRNAs were produced using overlapping nucleotides with the MegaScript T7 (Invitrogen/Thermo Fisher) *in vitro* transcription system. PCR templates for sgRNAs were produced using Phusion High-Fidelity DNA polymerase. The thermocycling conditions were as follows: 98 °C for 20 s; 35 cycles of 98 °C for 1 min s; 58 °C for 1 min; 72 °C for 1 min; and a final extension of 72°C for 7 min. Oligonucleotide sequences are given in Table 1. PCR products were purified using NucleoSpin Gel and PCR Clean-Up kit (Takara Bio, Kusatsu, Shiga, Japan), and 600 ng-1 ug of DNA templates were added to set up *in vitro* transcription reactions. Reactions were run for 16 hours at 37°C, treated with Turbo DNAse according to manufacturers’ instructions and purified using the MegaClear column purification kit (Thermo Fisher). The purified sgRNAs were tested with an *in vitro* cleavage assay. To produce a DNA template, genomic DNA (gDNA) from *Ae. aegypti* mosquitoes was purified using E.Z.N.A. MicroElute Genomic DNA Kit (Omega Bio-tek, Norcross, GA, USA) and the target region was amplified using Phire Animal Tissue Direct PCR Kit (Thermo Fisher) as described below. Reactions containing DNA template, individual sgRNAs and Cas9 protein in 1X NEB 3.1 buffer (New England Biolabs, Ipswich, MA, USA) were incubated at 37°C for 2 h, and diagnostic bands were visualized by electrophoresis on 1% agarose gel. sgRNAs for *Ae. aegypti* embryo injections were used at concentrations ranging between 70 ng/uL-180 ug/uL.

### Embryo injections and establishment of knock-out (KO) mosquito lines

Four to 5 days after blood feeding, 5-10 mated females were placed into a *Drosophila* vial with damp cotton and filter paper and placed in the dark for 50 min to stimulate oviposition. To generate heritable mutations in *Ae. aegypti* mosquitoes, mixtures of selected sgRNAs were injected into pre-blastoderm-stage embryos of Cas9-expressing mosquitoes 1-2 hours after laying. Briefly, embryos were aligned (with posterior poles on one side) on damp filter paper using a paintbrush, transferred on a glass slide using double-sided scotch tape, dried for 1 min and covered with a mixture of Halocarbon 700 oil and Halocarbon 27 oil (1:1) to prevent further desiccation. Embryos were injected into the posterior poles with quartz needles (QF100-70-10, Sutter Instrument, Novato, CA, USA) pulled by a Sutter P2000 needle puller (program 50, HEAT=500, FIL=5, VEL=50, DEL=128, PUL=0), using a Femtojet injector (Eppendorf, Hamburg, Germany) and an InjectMan micromanipulator using the following settings: injection pressure (pi) 1,000 hPa, compensation pressure (pc) 700 hPa, injection time (manual mode) 2-3 sec. After injection, embryos were transferred on the damp filter paper into egg cups with wet cotton, kept in the humid insectary for 4-5 days, and then hatched. Injected embryos (G_0_) that hatched and survived until adulthood were crossed individually. Legs of G_1_ mosquitoes were individually screened by PCR for the presence of deletions in the target gene as described below. A single heterozygous founder mosquito was outcrossed with wild-type age-matched *Ae. aegypti* mosquitoes to establish a KO mosquito line (see Results). As the target gene was located in the chromosome 1, the mutation was sex-linked [19]. As a result, to obtain homozygous mutants of both sexes, the selection process relied on chromosome recombination and identification of recombinant mosquitoes.

### Mosquito screenings for mutations

To screen live mosquitoes for the presence of deletions in the target gene, Phire Animal Tissue Direct PCR Kit (Thermo Fisher) was used according to manufacturers’ instructions. Briefly, mosquitoes were anesthetized on ice, a leg from each mosquito was removed using sharp forceps and immersed into 20 ul of sample dilution buffer supplemented with 0.5 ul of DNA release reagent. Leg samples in dilution buffer were incubated for 3 min at 98°C then used in PCR reactions. Primer sequences are provided in Table 1.

To characterize the αMan2 mutation at the transcript level, total RNA from αMan2 KO and wild-type mosquitoes was purified using E.Z.N.A. Total RNA kit (Omega Bio-tek) and cDNA synthesized using a gene-specific reverse primer and SuperScript III First-Strand Synthesis System (Thermo Fisher) according to manufacturers’ instructions. For cDNA synthesis, negative control gDNA from KO and wild-type mosquitoes was purified using E.Z.N.A MicroElute Genomic DNA Kit (Omega Bio-tek). PCR reactions were performed using Phire Animal Tissue Direct PCR Kit as described above. Information on primer sequences is in Table 1. For the detection of deletions in target genes, PCR products were separated by 2% agarose gel electrophoresis. Samples that separated into multiple bands were considered likely to contain an indel. The presence of αMan2 deletion(s) in both DNA and mRNA was then confirmed by PCR and direct sequencing of the target region.

### Quantification of relative Wolbachia density

Total DNA was extracted from *Wolbachia-* infected mosquito homogenates using a E.Z.N.A. Tissue DNA Kit (Omega Bio-Tek) kit according to the manufacturer’s instructions. qPCR was performed using PerfeCTa SYBR Green FastMix (Quantabio, Beverly, MA, USA) on a Rotor-Gene Q qPCR machine (Qiagen, Hilden, Germany) under the following thermocycling conditions: 95 °C for 2 min for initial denaturation; 40 cycles at 95 °C for 10 s, 60 °C for 40 s, 72 °C for 30 s for DNA amplification and data acquisition; 55–99 °C (5 s per increment) for the melt curve analysis. Relative *Wolbachia* densities were obtained by normalizing *Wolbachia* titers to the RpS17 gene levels as described previously [6]. Primer sequences are provided in Table 1. Crosses to introgress KO mutations into the *Wolbachia*-infected background are described in Results.

### Vector competence studies

Four-to-five-day old female mosquitoes were blood fed for approximately 1 hour on infected human blood containing 10^7^ infectious MAYV particles per mL or 10^5^ – 10^6^ infectious DENV particles per mL. After blood feeding, mosquitoes were anesthetized on ice and fully engorged females were transferred into cardboard cages; unfed females were discarded. Seven days post infection, mosquitoes were anesthetized using triethylamine (Sigma-Aldrich) and processed for vector competence assays. Mosquitoes were forced to salivate for 30 min into glass capillaries filled with a mix of 50% sucrose solution and FBS (1:1) to collect saliva samples. Body, legs, and saliva were then separately immersed in diluent solution containing 10% of FBS, 100 ug/mL of streptomycin, 100 units/mL of penicillin, 50 ug/mL gentamicin, and 2.5 μg/mL Amphotericin B in PBS. Body and legs samples were further homogenized by a single zinc-plated, steel, 4.5 mm bead using TissueLyser II (Qiagen) at 30 Hz for 2 min and centrifuged at 3,500 rpm at 4°C for 7 min in a bench top centrifuge to clear the homogenates. Samples were stored at −80°C. Virus titers in collected samples were determined by FFAs.

### Focus-forming assay for the quantification of MAYV and DENV

Vero or C6/36 cells were seeded in 96-well plates at a density of 3×10^4^ cells/well or 3×10^5^ cells/well for the titration of MAYV or DENV, respectively. Ten-fold serial dilutions (in serum-free medium) of virus samples obtained from mosquito bodies and legs were prepared and 30 uL of each were used in assays. Saliva samples were not further diluted. Cells were infected for 1 hour at 37°C or 28°C for MAYV and DENV assays, respectively. Infectious solutions were then removed and cells covered with 100 uL of complete growth medium (DMEM or RPMI) containing 0.8% methylcellulose (Sigma-Aldrich) and incubated at their respective temperatures. After 24 hours for MAYV or 3 days for DENV assays, overlay medium was removed, cells were fixed with 4% formalin (Sigma-Aldrich) in PBS (Gibco/Thermo Fisher) for 15 min and permeabilized with 0.2% TritonX in PBS for 15 min. Primary antibodies were diluted in PBS and incubated overnight at 4°C. Secondary antibodies were incubated overnight at 4°C for MAYV assays and 3 hours at room temperature for DENV. After the final wash, cells were dried briefly, and MAYV or DENV foci immediately counted using an Olympus BX41 inverted microscope equipped with an UPlanFI 4X objective and a FITC filter.

### Data analysis

Infection, dissemination, and transmission rates were analyzed using contingency tables. Data on *Wolbachia* titers were analyzed by Mann-Whitney U tests. Due to violation of the equal variance assumption, data on viral titers were analyzed using the Brown, Forsythe ANOVA method with Welch’s correction for multiple tests.

## Results

### Generation of Wolbachia-negative and Wolbachia-positive αMan2 KO Ae. aegypti mosquitoes

To generate a deletion in the αMan2 gene, *Ae. aegypti* embryos expressing Cas9 protein (G_0_, N=115) were injected with a mix of four sgRNAs targeting exon 5 of the gene. Surviving G_0_ individuals (females N=19, males N=10) were outcrossed to wild-type mates (1 male per 1-2 females), females blood fed, eggs collected, and hatched in small batches for further screening (Fig. 1A). Forty G_1_ male mosquitoes were individually screened by PCR for the presence of deletions in the target gene. Three G_1_ males with αMan2 deletions were identified: two with an identical 13 nt deletion and one with a double deletion allele consisting of a 46 nt deletion at one sgRNA target site and a 9 nt deletion at another sgRNA target site (55 nt deletion total) (Supplementary Figure 1A). This 55 nt deletion was predicted to result in a 155-amino acid-long truncated protein instead of a 1174-amino acid long functional enzyme. The male mosquito with two deletions (totaling 55 nt) was further crossed with 7 age-matched wild-type females to establish a line (Fig. 1A).

**Figure 1.**
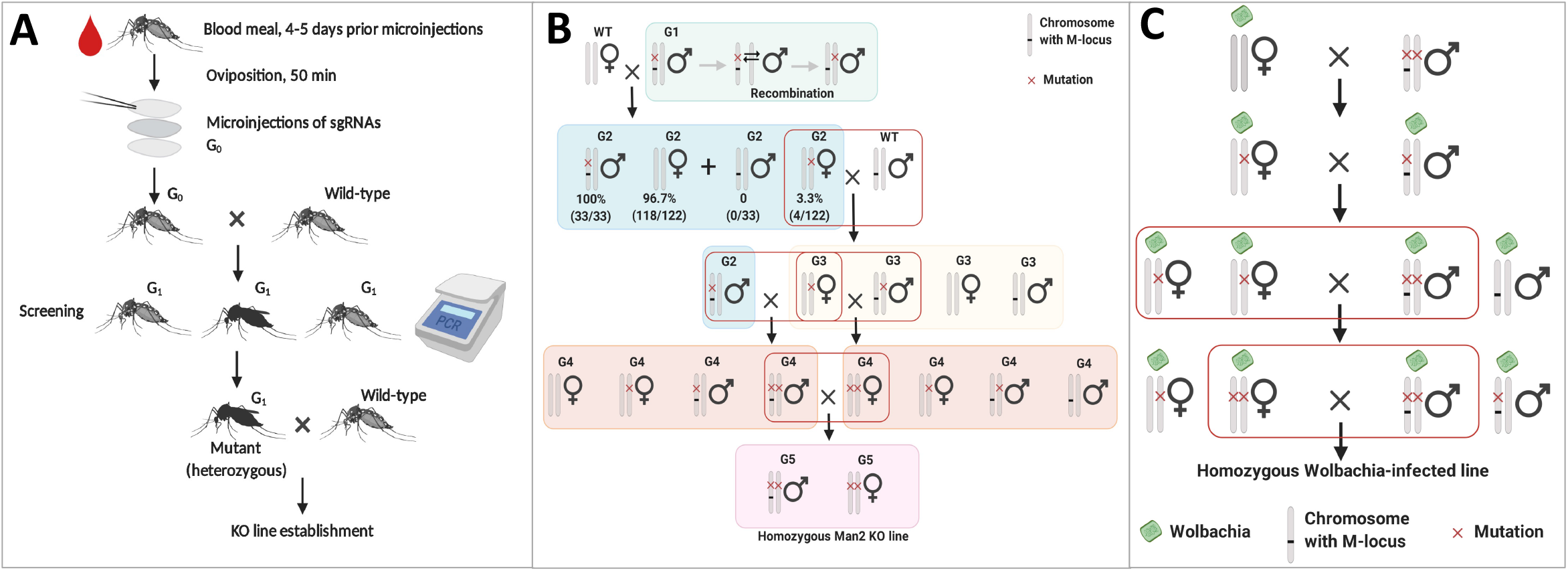
Overview of the approach used to generate *Ae. aegypti* strains used in this study. A) Generation of **a**Man2 KO mutations using CRISPR-Cas9 mutagenesis. B) Crossing scheme to generate a homozygous **a**Man2 KO line. C) Crossing scheme to generate a homozygous **a**Man2 KO line infected with *Wolbachia*.

Since αMan2 is located in on chromosome 1, deletions in this gene were expected to be sex-linked [19]. All G_2_ male progeny from the selected heterozygous G_1_ mutant mosquito that were screened (N=33) carried deletions, while the majority of G_2_ females were wild-type. To obtain mutant females, we relied on identification of recombinant mosquitoes. Four out of 122 screened G_2_ females (3.3%) were recombinants and carried a deleted copy of αMan2. These G_2_ females were further crossed with wild-type males to obtain G_3_ males with the mutation on the opposite chromosome. Homozygous mutant males were obtained via crossing G_3_ mutant females and the generated G_2_ mutant males. Homozygous mutant males and females were selected and crossed to obtain a homozygous αMan2 KO line (Fig. 1B). The presence of αMan2 deletions in both DNA and mRNA was confirmed by PCR and direct sequencing of the target region (Supplementary Figure 1B).

To generate a *Wolbachia*-infected αMan2 KO line, *Wolbachia*-infected *Ae. aegypti* females we crossed with αMan2 KO *Wolbachia*-negative male mosquitoes so that CI would not sterilize the cross [4]. Every generation after crossing was checked by PCR for the presence of both the mutation and *Wolbachia* infection, and heterozygous *Wolbachia*-infected males and females were crossed. Homozygous *Wolbachia*-infected males and heterozygous *Wolbachia*-infected females were selected and further crossed as described above to obtain homozygous *Wolbachia*-infected male and female mosquitoes. Homozygous *Wolbachia*-infected male and female mosquitoes were selected using PCR and crossed to establish a pure homozygous *Wolbachia*-infected αMan2 KO mosquito line (Fig. 1C). *Wolbachia*-negative male mosquitoes from the parental Cas9-expressing *Ae. aegypti* line, which was used for embryo injections, were crossed with *Wolbachia*-infected *Ae. aegypti* females following similar procedure as described above to obtain a *Wolbachia*-infected wild-type control line with comparable genetic background for infection experiments.

### *Effect of αMan2 KO on Wolbachia titers in Ae. aegypti* mosquitoes

We tested the effect of gene KO on *Wolbachia* titers by qPCR and found that the αMan2 KO mutation significantly increased mean *Wolbachia* levels by approximately 2-fold compared to the wild-type genetic background (Fig. 2).

**Figure 2.**
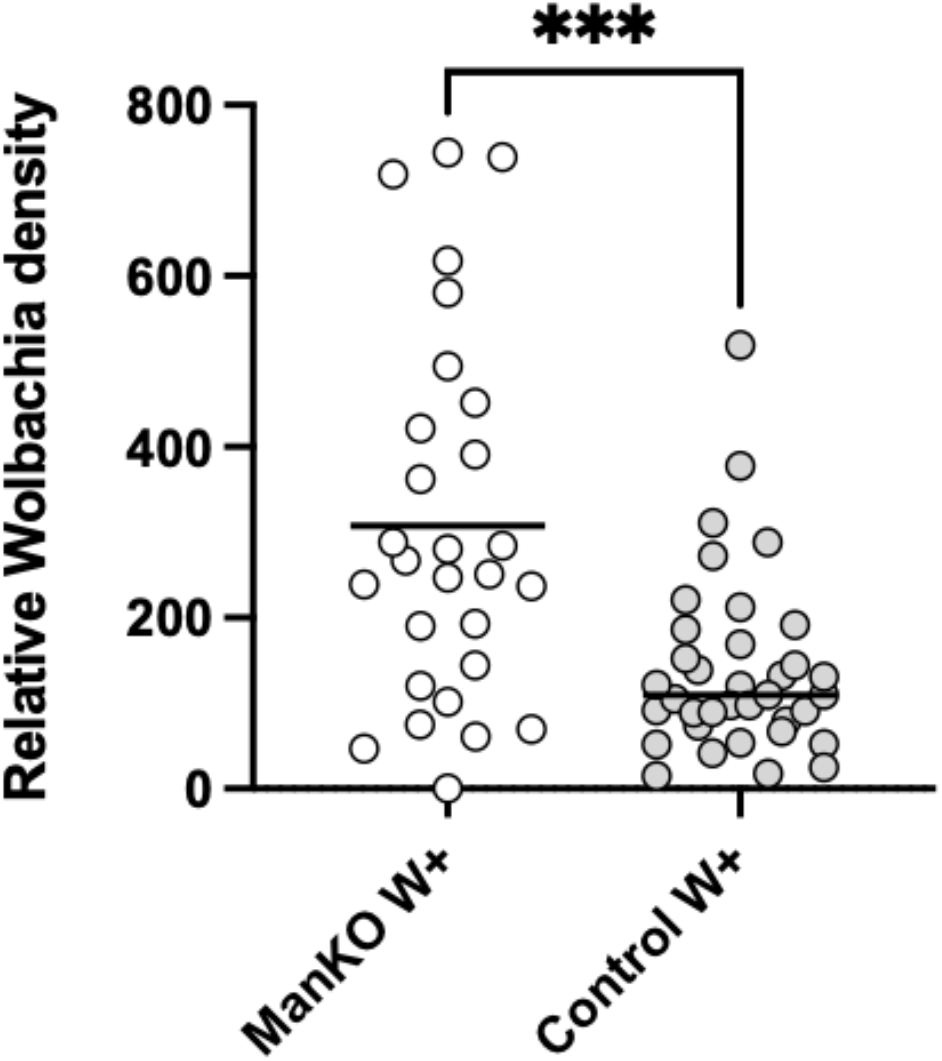
*Wolbachia* titers in **a**Man2 KO and wild-type mosquitoes. Mutant mosquitoes had significantly higher levels of *Wolbachia* compared to wild-type (*P* = 0.0007).

### Effect of Wolbachia and αMan2 KO on DENV infection, dissemination, and transmission rates

DENV infection and dissemination rates were lower, but not significantly so, in *Wolbachia*-infected wild-type mosquitoes (50% vs. 36% infection). In *Wolbachia*-uninfected mosquitoes, the αMan2 KO mutation was associated with significantly reduced DENV infection rates (16% vs 50%). Although either *Wolbachia* alone or the αMan2 KO mutation alone both tended to reduce DENV infection and dissemination rates, the two effects were not additive. Rather, there was an interaction between *Wolbachia* infection status and genotype; when the αMan2 KO mutation was present in a *Wolbachia*-infected background, infection rates were similar to *Wolbachia*-uninfected wild-type mosquitoes (50% vs. 46%) (Table 2). We did not observe DENV transmission in any treatment.

**Table 2.** Virus infection, dissemination, and transmission rates for experimental treatments 7 days post-infection.

| Table 2. |  |  |  |  |  |  |  |  |  |
| --- | --- | --- | --- | --- | --- | --- | --- | --- | --- |
| Virus | Geneotype | Wolbachia | N | Infection % | Sig | Dissemination % | Sig | Transmission % | Sig |
| DENV | WT | No | 36 | 0.50 | a | 0.22 | a | 0.00 | a |
| DENV | WT | Yes | 69 | 0.36 | a | 0.07 | a,b | 0.00 | a |
| DENV | Man2 KO | No | 37 | 0.16 | b | 0.03 | a,b | 0.00 | a |
| DENV | Man2 KO | Yes | 46 | 0.46 | a | 0.02 | b,c | 0.00 | a |
| MAYV | WT | No | 71 | 1.00 | a | 1.00 | a | 0.42 | a |
| MAYV | WT | Yes | 64 | 0.45 | b | 0.19 | b | 0.00 | c |
| MAYV | Man2 KO | No | 43 | 0.98 | a | 0.95 | a | 0.33 | b,c |
| MAYV | Man2 KO | Yes | 66 | 0.27 | c | 0.08 | b | 0.00 | c |

### Effect of Wolbachia and αMan2 KO on DENV titers in mosquitoes

In a wild-type genetic background, DENV titers were significantly lower in *Wolbachia*-infected mosquitoes compared to uninfected (Fig. 3A); a demonstration of canonical *Wolbachia*-induced PB. In a *Wolbachia*-uninfected background, the αMan2 KO mutation itself reduced DENV titers (Fig. 3A). We observed an interaction between *Wolbachia* infection status and genotype, where the αMan2 KO mutation in a *Wolbachia*-infected background reduced the ability for *Wolbachia* to suppress DENV (Fig. 3A). DENV dissemination titers in mosquito legs between treatments did not significantly differ, possibly due to lower dissemination rates and resulting lack of power to detect a statistical difference (Supplementary Fig. 2A).

**Figure 3.**
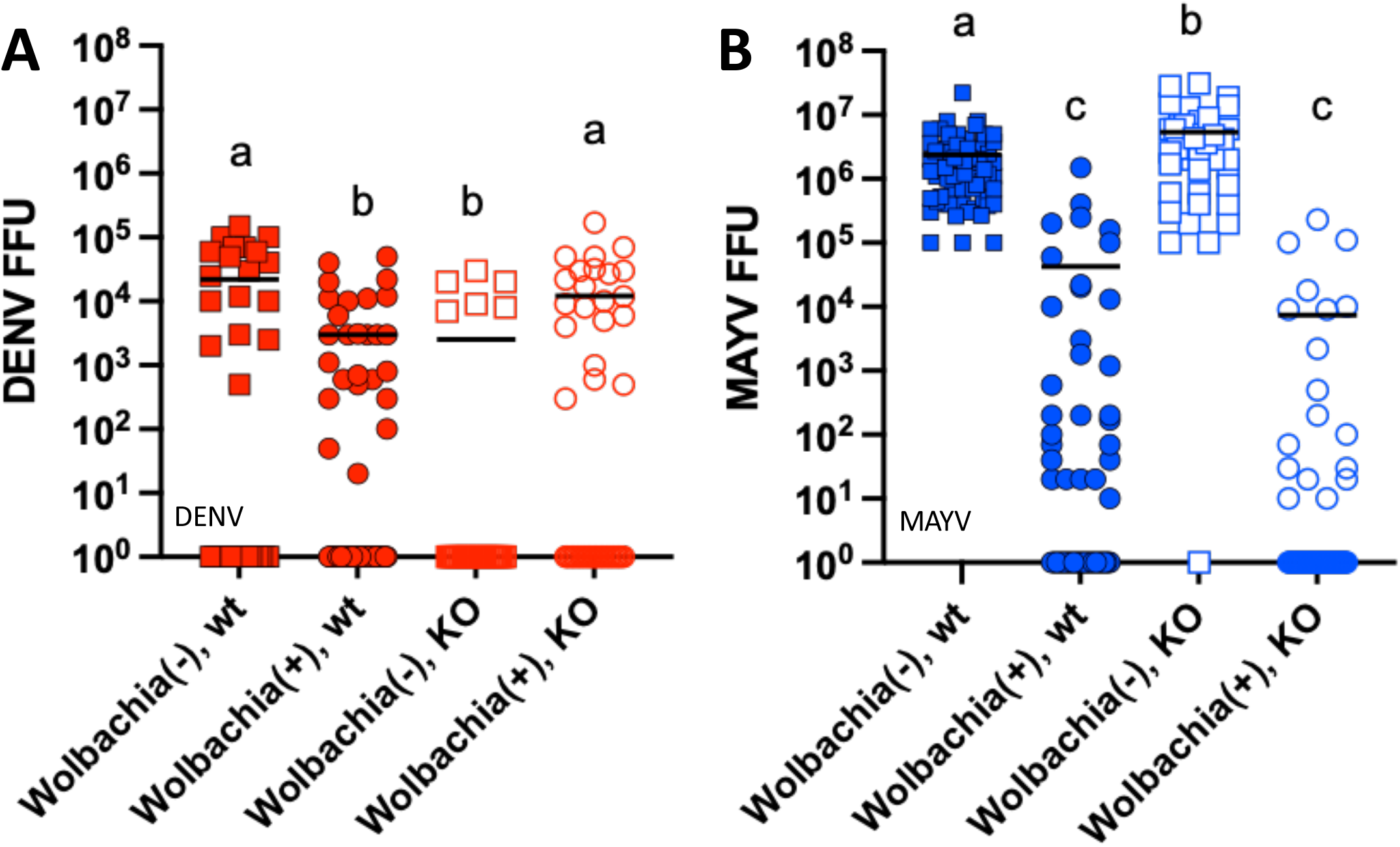
A) DENV) and B) MAYV body titers in experimental mosquitoes 7 days post-infection. Red = DENV; Blue = MAYV; Squares = *Wolbachia*-uninfected; Circles = *Wolbachia*-infected; Filled symbols = wild-type; open symbols = αMan2 KO. Viruses were analyzed separately; treatments with different letters are significantly different (*P* < 0.01).

### Effect of Wolbachia and αMan2 KO on MAYV infection, dissemination, and transmission rates

MAYV infection and dissemination rates were higher generally compared to DENV, perhaps due to higher initial viral titers. 100% of wild-type, *Wolbachia*-uninfected mosquitoes became infected with and disseminated MAYV. *Wolbachia*-infected, wild-type mosquitoes had significantly reduced infection (45%) and dissemination (19%) rates as would be expected from *Wolbachia*-induced PB. Infection and dissemination rates in *Wolbachia* uninfected αMan2 KO mosquitoes were similar to uninfected wild-type mosquitoes. αMan2 KO, *Wolbachia*-infected mosquitoes had the lowest infection (27%) and dissemination (8%) rates, opposite to what was observed for DENV. We did observe transmission of MAYV in these experiments, where the highest transmission rate (42%) was observed in wild-type *Wolbachia*-uninfected mosquitoes, and no transmission was observed in *Wolbachia*-infected mosquitoes, regardless of genotype (Table 2). *Wolbachia* uninfected KO mosquitoes had intermediate transmission rates (33%) (Table 2).

### Effect of Wolbachia and αMan2 KO on MAYV titers in mosquitoes

In a wild-type genetic background, MAYV titers were significantly lower in *Wolbachia*-infected mosquitoes compared to uninfected (Fig. 3B); again consistent with canonical *Wolbachia*-induced PB. However, in a *Wolbachia*-uninfected background, the αMan2 KO mutation was associated with enhanced MAYV titers compared to wild-type mosquitoes (Fig. 3B). In a *Wolbachia*-infected background, the αMan2 KO mutation did not affect PB, and MAYV titers were indistinguishable from *Wolbachia*-infected wild-type mosquitoes (Fig 3B). MAYV dissemination titers in mosquito legs between treatments significantly differed in a similar pattern to body titers (Supplementary Fig. 2B).

## Discussion

A recent genetic screen identified that single-nucleotide polymorphisms in the *Ae. aegypti* αMan2 gene were associated with stronger or weaker *Wolbachia* (wMel)-mediated PB of DENV [6], but the functional role of this gene in DENV blocking remains unclear. Due to the intronic location of the identified polymorphisms, it was hypothesized that they could affect gene expression or splicing; however, no significant differences in αMan2 expression were found between selected low and high blocking mosquito populations [6]. We recently published a study [20] using RNAi to knock down expression of αMan2 in *Wolbachia* infected and uninfected *Ae. aegypti* to examine its effect on PB for DENV and Chikungunya virus (CHIKV); an alphavirus closely related to MAYV [21]. RNAi demonstrated some influence of αMan2 in PB, but results were not dramatic [20]. While RNAi silencing has become a cornerstone of genetic analysis in mosquitoes, the effects of this manipulation are generally transient, with the length and strength of silencing varying depending on the tissue and the target gene [22]. In addition, protein turnover dynamics can affect the strength of any phenotypic effects resulting from gene knockdown [23]. Due to these issues, genetic ablation of the target gene of interest allows for much more robust interrogation of gene function. We used CRISPR-Cas9 gene editing to generate αMan2 KO mutations in *Ae. aegypti* mosquitoes to functionally investigate the role of this gene in arbovirus replication and found that the αMan2 KO mutation affected arboviruses in a pathogen and *Wolbachia* infection-specific manner. This is especially interesting as the *Wolbachia* strain used in the original genetic screen [6] was *w*Mel (originally from *Drosophila melanogaster*), while we performed experiments using the *Wolbachia* strain *w*AlbB (originally from *Ae. albopictus*). These two *Wolbachia* strains are not closely related, yet both seem to interact with αMan2, suggesting that candidate genes identified by Ford et al. [6] may be broadly applicable across different *Wolbachia* strains.

Differences in viral phenotypes between mutant and wild-type mosquitoes in a *Wolbachia*-infected background cannot be explained by a direct effect of the KO mutation on *Wolbachia* titers. *Wolbachia* levels were approximately twice as high in αMan2 KO mosquitoes compared to wild-type. While *Wolbachia*-induced suppression of MAYV was similar in both αMan2 KO and wild-type mosquitoes, DENV was not blocked in *Wolbachia*-infected mutant mosquitoes, highlighting the complex interactions between the mosquito genome, *Wolbachia*, and the specific viral pathogen. *Wolbachia* loads can be affected by mosquito immunity [24], and the mosquito immune system can be modulated by glycosylation pathways [25], suggesting a potential explanation for higher *Wolbachia* titers in αMan2 KO mosquitoes, although this phenomenon requires further study.

For DENV, the αMan2 KO mutation itself conferred some resistance to virus, significantly reducing viral titer. *Wolbachia* alone also reduced viral titer. However, there was an interaction between αMan2 genotype and *Wolbachia* infection; when the mutation was coupled with *Wolbachia* in the mosquito, DENV infections were no longer suppressed. We observed a different phenomenon with MAYV. In a *Wolbachia*-uninfected background, the αMan2 KO mutation did not significantly alter viral infection rates but did significantly enhance viral titers in the mosquito. In a *Wolbachia*-infected background, the mutation increased the ability for *Wolbachia* to suppress viral infection rates and did not interfere with the ability for *Wolbachia* to suppress viral titers, although it did not further enhance *Wolbachia* PB.

The fact that the αMan2 mutation (in the absence of *Wolbachia*) had different effects on DENV vs. MAYV is not necessarily surprising, as DENV is a flavivirus, while MAYV is an alphavirus. These two viral families are not closely related, and it has been demonstrated that the mosquito immune system responds differently to diverse viral groups [26]. The fact that the αMan2 KO mutation can have different effects on how *Wolbachia* suppresses different viral families is perhaps also not surprising, as *Wolbachia* has been shown to differentially suppress different pathogens in other systems [27–29]. Ultimately, these data demonstrate the complexity of the *Wolbachia* PB phenotype. In their screens, Ford et al. [6,7] identified dozens of potential candidate genes regulating PB; here we have disrupted one of them. It is likely that disruption of other candidates could have equally complex consequences, to say nothing of multiple stacked mutations.

While our data show that *Ae. aegypti* αMan2 is a modulator of arbovirus infection, and involved in the *Wolbachia* PB phenotype, the mechanism by which it works, and has variable effects on different viruses, remains unclear. αMan2 is involved in protein glycosylation [8], which may affect viral biogenesis, replication, and infectivity [11], so it is logical that disruption of this gene would affect viral infection phenotypes. However, CRISPR is a blunt tool, and further molecular research is necessary to determine the specific mechanism by which αMan2 modulates replication of specific viruses and how it contributes to *Wolbachia* PB.

## List of abbreviations

cDNA: complementary DNA
CI: cytoplasmic incompatibility
DENV: dengue virus
DMEM: Dulbecco’s Modified Eagle Medium
DPI: days post infection
FBS: Fetal bovine serum
FFA: focus-forming assay
FFU: focus-forming unit
gDNA: genomic DNA
GE: Genome equivalent
KO: knockout
MAYV: Mayaro virus
PB: pathogen blocking
PBS: Phosphate-buffered saline
PFU: Plaque-forming unit
qPCR: Quantitative polymerase chain reaction

## Declarations

### Ethics approval and consent to participate

Not applicable.

### Consent for publication

Not applicable.

### Availability of data and materials

All data generated or analyzed during this study are included in this published article.

### Authors’ contributions

NU, VMM, EAM and JLR conceived and designed the study; NU, REJ, AH, VMM, MJJ, LTS performed the experiments; NU, JLR analyzed the data; OSA, ML, ZX, KL, RS, EAM, JLR contributed reagents and materials; NU, JLR drafted the manuscript; all authors read and approved the final version of the manuscript.

## Acknowledgments

We would like to thank Dr. Duverney Chaverra-Rodriguez for valuable advice concerning CRISPR experiments and Hillery Metz for valuable comments on a draft version of the manuscript.

---

**Supplementary Figure 1.**
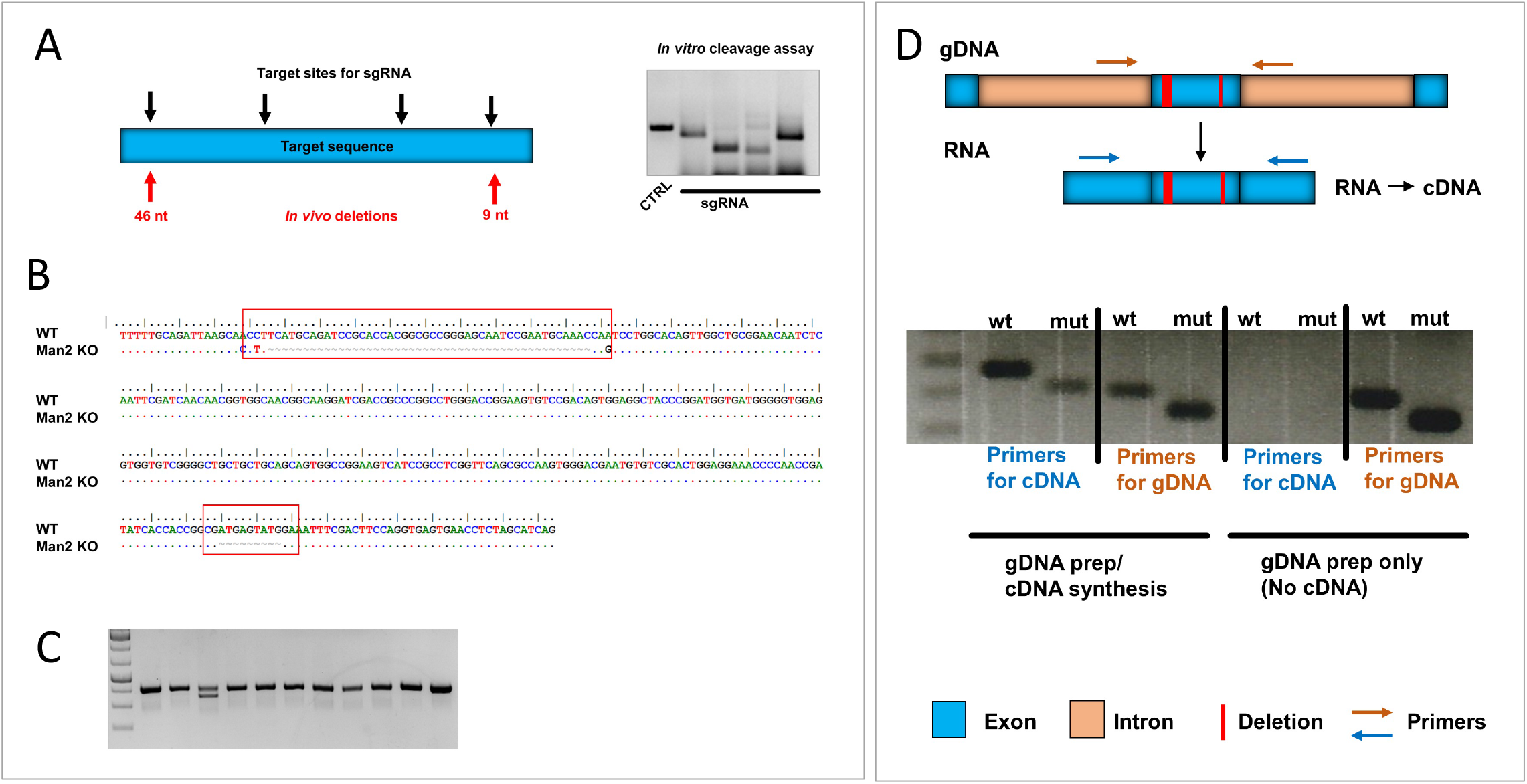
Verification of gene deletions. A) *In vitro* cleavage assay to validate sgRNAs; B) Sequencing data for identified male mosquito with two deletions totaling 55 nt in the αMan2 gene: colored dots represent identical nucleotides, grey tildes represent deleted nucleotides; C) PCR identification of heterozygous male mosquito carrying the αMan2 55 nt mutation; D) Detection of αMan2 deletion in gDNA and mRNA. Two sets of primers were used: one that binds gDNA in intronic regions outside the target exon and one that binds cDNA in adjacent exons. The former primer set does not yield any product on the cDNA template as the binding regions are removed during RNA splicing. The latter primer set does not yield any product under the given PCR conditions if only gDNA template is present, as in gDNA, primers are separated by extended intronic regions and the fragment doesn’t amplify.

**Supplementary Figure 2.**
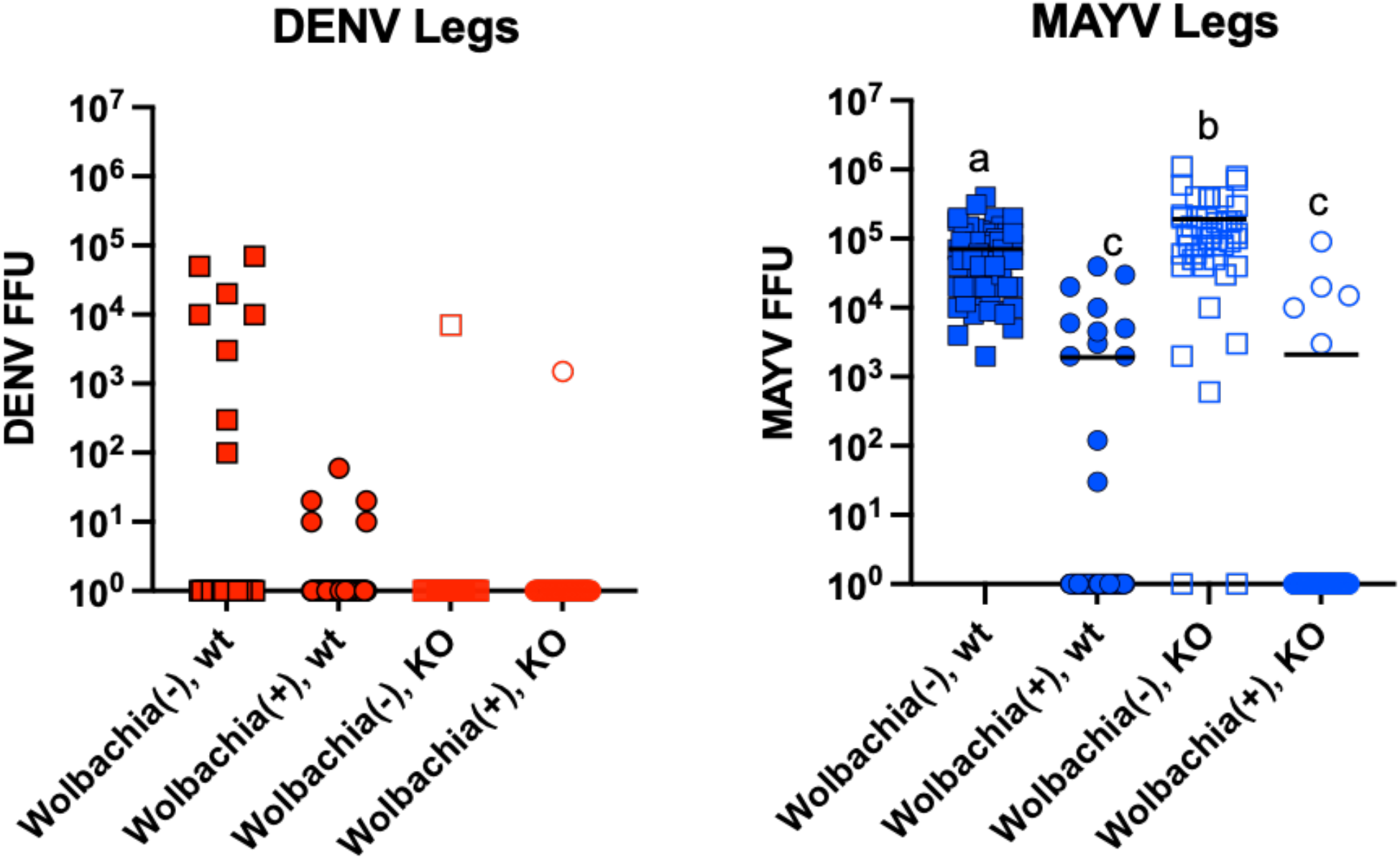
A) DENV) and B) MAYV leg titers in experimental mosquitoes 7 days post-infection. Red = DENV; Blue = MAYV; Squares = *Wolbachia*-uninfected; Circles = *Wolbachia*-infected; Filled symbols = wild-type; open symbols = αMan2 KO. Viruses were analyzed separately; treatments with different letters are significantly different (*P* < 0.001).

